# The duration of caffeine treatment plays an essential role in its effect on sleep and circadian rhythm

**DOI:** 10.1101/2021.10.08.463733

**Authors:** Aishwarya Segu, Nisha N Kannan

## Abstract

Sleep is regulated by the homeostatic system and the circadian clock. The circadian clock governs the timing of sleep-wake cycles and the homeostatic mechanisms modulate the amount and depth of sleep. Caffeine intake promotes wakefulness in *Drosophila*. In humans, caffeine is consumed on a daily basis and hence it is important to understand the effect of prolonged caffeine intake on both circadian and homeostatic regulation of sleep. Furthermore, sleep changes with age and the impact of caffeine on age dependent sleep fragmentation is yet to be understood. Hence in the present study, we examined the effect of short exposure to caffeine on homeostatic sleep and age dependent sleep fragmentation in *Drosophila*. We further assessed the effect of prolonged exposure to caffeine on homeostatic sleep and circadian clock in *Drosophila*. The results of our study showed that short exposure to caffeine reduces sleep in flies and enhances sleep fragmentation with increasing age. On the other hand, prolonged caffeine exposure did not exert any significant effect on duration of sleep in mature flies. Nevertheless, prolonged caffeine ingestion decreased the morning and evening anticipatory activity in these flies indicating that it affects the circadian rhythm. These flies also exhibited phase delay in the clock gene *timeless* transcript oscillation and altered behavioral rhythm with either a longer free running period or arrhythmicity under constant darkness. In summary, the results of our studies showed that short exposure to caffeine increases the sleep fragmentation with age whereas prolonged caffeine exposure disrupts the circadian clock.

**Statement of Significance:** In the present study, we assessed the effect of duration of caffeine intake on age dependent sleep fragmentation and circadian clock by using *Drosophila* as a model organism. We found that short exposure to caffeine reduced sleep and food intake in flies. It also increased the sleep fragmentation with age. On the other hand prolonged caffeine exposure did not affect the sleep and feeding indicating that these flies developed tolerance to caffeine. As in mammals, caffeine slows down the circadian clock in flies. Most importantly, prolonged caffeine treatment disrupted the circadian rhythm in *Drosophila*. Our studies provide new insights into the differential effect of duration of caffeine intake on circadian clock and sleep.

## Introduction

Sleep is a well-documented and conserved behavior across kingdoms^1^. Sleep is critical for diverse physiological functions like metabolism, immune response^2–4^, cognition and memory formation^5^. Sleep is majorly regulated by two processes namely the circadian clock that regulates the timing of sleep^6^ and the homeostatic pathways that modulates the intensity and depth of sleep^1^. This regulation is best understood by the two-process sleep model proposed by Borbeley, in 1981^7^. This model has been successful in simulating the sleep timing and depth in various experimental conditions^7^. Accumulating evidence continues to help us understand the interactions between these two processes. Electrophysiological recordings in the suprachiasmatic nucleus (SCN), the central pacemaker in mammals, reported continuous interaction between the circadian clock and sleep homeostat indicating that the interplay between these two systems is important in governing sleep^7^.

The circadian clock that controls the timing of sleep consists of interlocked transcriptional-translational feedback loop (TTFL) composed of *period* (*per*) and *timeless* (*tim*) genes which is activated when the positive transcription factors CLOCK (CLK) and CYCLE (CYC) bind to their E-box promoters^8,9^. This transcriptional - translational feedback loop maintains the transcript and protein oscillations that govern the circadian rhythm in *Drosophila*^10^. Studies in the recent past have unraveled that circadian neuropeptide - pigment-dispersing factor (PDF); diuretic hormone 31 (DH31), glutamate and GABA_A_ receptor *Resistant to dieldrin* (RDL) expressed in subsets of circadian clock neurons mediate sleep and wakefulness to occur at the specific time of the day^11–13^. Additionally, circadian output molecule WIDE AWAKE (WAKE) upregulates GABA_A_ receptor RDL and reduces the excitability of clock neurons at the day/night transition to promote sleep onset.

The sleep homeostat controls the depth and intensity of sleep^14–16^. Sleep depth is highly variable depending on multiple environmental factors and sleep pressure^17^. One of the major aspects that decides the sleep pressure is the prior wakefulness period. An elevation in sleep pressure accounts for the immediate increase in sleep, termed as sleep rebound observed after a period of sleep deprivation. Sleep loss also increases the overall abundance of synaptic proteins and impairs the neuronal homeostasis^18,19^. From behavioral, neurogenetic and physiological approaches, researchers have dissected the neural circuits involved in homeostatic regulation of sleep. In *Drosophila*, a cluster of neurons projecting into the dorsal fan-shaped body (dFSB) present in the central brain complex regulates sleep homeostasis^20,21^. Along with the dFSB, mushroom body (MB) and R2 neurons in the ellipsoid body (EB) are the important brain centers mediating sleep drive and arousal in *Drosophila*^17,22^.

Caffeine, a psycho-stimulant used widely for promoting wakefulness, is an antioxidant derived from plant based sources. In *Drosophila,* caffeine is shown to increase calcium signaling in the dopaminergic neuron (DAN’s) sub-clusters called the protocerebral anterior medial (PAM) neurons^23^. Further, it was also shown that activity of caffeine required dopaminergic receptor 1 (DopR1) expressed in MBs to cause increased wakefulness in flies^24^. The increased wakefulness observed due to caffeine treatment leads to decreased sleep in flies. As sleep is governed by both the circadian clock and homeostatic system, it is important to understand the effect of caffeine on both of these pathways.

In *Drosophila*, development influences sleep during the early adult phase leading to progressive decrease in sleep from young to mature flies and further, sleep quality decreases with increasing age^17,25,26^. Previously it has been shown that caffeine affects sleep duration and also causes sleep fragmentation^23^. But, as sleep quality is age dependent, the effect of caffeine on sleep may vary with increasing age. Therefore, in our present study we assessed the effect of caffeine on age dependent sleep fragmentation in *Drosophila*. The results of our study showed that short exposure to caffeine reduces sleep in young and old flies. It also increased the age dependent sleep fragmentation. Furthermore, we also addressed the effect of prolonged caffeine exposure on sleep and circadian clock in *Drosophila*. Prolonged exposure to caffeine did not reduce the sleep whereas it delayed the phase of core-clock gene *timeless* transcript oscillation and also resulted in either longer free running period or behavioral arrhythmicity in mature flies. Furthermore prolonged caffeine treatment delayed the pre adult development and reduced the life span in *Drosophila*.

## Materials and methodology

### Fly stock and maintenance

*w^1118^* flies were obtained from the Bloomington Drosophila Stock Centre (BDSC #5905). All the flies were raised in cornmeal dextrose medium and were maintained under 12 hour (h) light : 12 h dark (LD) cycle where lights came on at Zeitgeber Time 00 (ZT 00) and went off at ZT12 in the *Drosophila* growth chamber (Percival Scientific, Perry IA) at 25° C temperature with 65 ± 5% humidity.

### Short exposure to caffeine

To record the sleep under short exposure to caffeine, 1, 10, 20 and 30 day old *w^1118^* male flies were transferred to locomotor activity glass tubes (65 mm length, 5 mm diameter) containing cornmeal dextrose medium for acclimatization 12h prior to the start of the experiment. These flies were transferred into locomotor activity glass tubes containing cornmeal dextrose medium with different concentrations of caffeine (0.0, 0.5, 0.75 and 1 mg/mL) just before ZT 00 on the day of locomotor activity recording. Sleep was recorded by using the Drosophila Activity Monitors (DAM, Trikinetics, USA) for 24h under LD in a cooled incubator (MIR-154, Panasonic, Japan). On the subsequent day, at ZT 00, these flies were transferred back to fresh cornmeal dextrose medium without caffeine, to record their sleep rebound for the next 24h. Activity counts were measured at every 1 minute (min) interval and sleep was defined as 5 min or more of continuous inactivity. Sleep parameters such as total sleep duration, sleep bout length, sleep bout number, sleep latency and sleep in 1h were analyzed using Sleep and Circadian Analysis MATLAB Program (SCAMP)^27^. To confirm the results, each experiment was replicated thrice with a sample size of 25-32 flies in each replicate. The data obtained from one such biological replicate is used in the results and figures.

### Prolonged caffeine exposure

For prolonged caffeine exposure, we reared first instar larvae in cornmeal dextrose medium containing 0.5, 0.75 and 1 mg/mL of caffeine. Freshly emerged *w^1118^* male flies were also maintained in vials with the respective caffeine containing medium. To assess the effect of prolonged caffeine exposure on sleep, 10 day old mature flies under prolonged caffeine treatment were transferred into the locomotor activity glass tubes containing the cornmeal dextrose medium with the respective caffeine concentrations. Sleep for 24h and sleep rebound was recorded as mentioned in short exposure to caffeine.

Further, to assess the effect of prolonged caffeine exposure on the circadian clock, 10 day old male flies under prolonged caffeine caffeine treatment were individually introduced into locomotor activity glass tubes containing standard cornmeal medium with different concentrations of caffeine. To assess the effect of caffeine on clock mediated activity-rest rhythm, locomotor activity was recorded under LD with caffeine for the first 5 days. Subsequently their locomotor activity was recorded under constant darkness (DD) with caffeine for 10 days to assess the free running period. The normalized waveform of activity-rest rhythms under LD was obtained by dividing the activity data collected in every 1h interval by the total amount of activity during the 24h multiplied by 100. Anticipation index (AI) for lights-on and lights-off was calculated as the ratio of activity in 3h just prior to lights-on or lights-off to the activity that occurs 6h before the transition^28^. The free running period of activity-rest rhythm was estimated by using the Lomb Scargle Periodogram of CLOCKLAB, Actimetrics, USA. For rhythmicity analysis, autocorrelation periodograms were constructed using raw activity count data and rhythmicity index (RI) was calculated using VANESSA^29^. Flies were considered rhythmic if RI<0.3, weakly rhythmic if 0.01>RI>0.3. Each experiment was repeated thrice with 28-32 flies in each replicate. One of such biological replicates is shown in the sleep and locomotor activity plots under LD. Whereas, a sample size of 80-90 flies was used for assessing the free running period.

### Adult stage specific caffeine treatment

For the adult stage specific caffeine treatment, freshly emerged *w^1118^* male flies raised in cornmeal medium were transferred and maintained in vials containing cornmeal medium with different caffeine concentrations (as mentioned above). After 10 days of adult stage specific caffeine treatment, sleep, sleep rebound, activity-rest rhythm under LD and DD was recorded by using the protocol mentioned in prolonged caffeine exposure.

### CApillary FEeder assay (CAFE)

To study the feeding behavior, we used a modified CApillary FEeder assay (CAFE)^30^ using microtips (Details of modified CAFE protocol is available in Supplementary S4). To measure food intake, 16 male flies were introduced into CAFE assay tubes and the food containing 0.1M sucrose solution along with 1% Orange-G (Sigma Aldrich) solution was provided. Three such replicates each containing 16 flies were used in this study. To test the effect of short exposure to caffeine on feeding, after 24h of caffeine treatment, the flies were introduced into CAFE assay tubes containing different concentrations of caffeine along with sucrose/Orange-G solution during the feeding assay. To assess the food intake post caffeine treatment, flies kept under caffeine containing medium for 24h were removed and introduced into CAFE tubes containing only 0.1M sucrose solution with 1% Orange-G without caffeine during the feeding assay. The feeding was recorded for 3h starting from ZT 01 and the volume of food intake in each CAFE tube was measured using vernier calipers. Further, the food intake of each fly was calculated by dividing the volume of food intake in each CAFE tube with the number of flies present in that tube. To assess the effect of prolonged and adult-stage specific caffeine treatment on feeding, the food intake was assessed in 10 day old flies under prolonged and adult-stage specific caffeine treatment respectively (by using the same protocol mentioned for short term exposure to caffeine).

### Total RNA isolation and quantitative real time PCR

To assess the effect of prolonged caffeine treatment on clock gene transcript oscillation, 10 day old male flies entrained to LD under cornmeal dextrose medium and under 1 mg/ml of prolonged caffeine treatment were used. Three biological replicates each consisting of 30 male flies were flash frozen using liquid nitrogen with 4h time intervals from ZT 02 to 22 under LD. To check the expression of genes involved in homeostatic sleep regulation we sampled the flies only at one time point - ZT 14. The heads were separated and total RNA isolation was performed using TRIzol (Invitrogen) Phenol:Chloroform (Sigma aldrich) method. The heads were crushed in TRIzol and post incubation was separated into aqueous layer using 99.5% chloroform. The aqueous layer was precipitated using Isopropanol (Sigma Aldrich). The RNA was quantified using nanodrop and cDNA was synthesized using Takara PrimeScript 1st strand cDNA synthesis kit (Cat #6110A) using the manufacturer’s protocol. Further quantitative real time PCR was performed using Takara TB green Premix Ex Taq II intercalating agent (Cat #RR820A) and detected using Bio Rad CFX96 Touch Real-Time PCR. The *timeless* mRNA values are shown after normalization with the house-keeping gene *rp49*. The primer details are provided in table S1.

### Jonckheere-Terpstra-Kendall (JTK) analysis for checking gene oscillation

To analyze *timeless* transcript oscillation we used the JTK algorithm from Metacycle^31^ using R-progamming. The mean from technical replicates of each biological replicate was used for analysing the transcript oscillation. The transcript levels were considered to be oscillating if the *p*-value was less than 0.05. For transcripts which showed oscillation, the phase was calculated.

### Life span assay

To test the effect of prolonged caffeine treatment on life span, larvae were reared under different caffeine concentrations and freshly emerged 10 male flies were transferred into vials containing cornmeal dextrose medium with 0.5, 0.75 and 1 mg/mL of caffeine. Ten such replicate vials were set up for each caffeine concentration under LD in a cooled incubator (Percival Scientific, Perry IA). Flies were provided with fresh medium on every third day to avoid death due to desiccation. The death occurred on each day was noted until all flies died and survival curves were analyzed using a log-rank test.

### Pupation time assay

50 first instar larvae were transferred into the vial containing either non-caffeinated or caffeinated food. Each biological replicate consisting of three such technical replicates were maintained under LD in a cooled incubator (Percival Scientific, Perry IA). Their pupation was assessed after the third instar larval stage with 6h interval starting from ZT 00 to ZT 18. Average of three biological replicates were used to quantify the pupation time. The percentage of pupation was analyzed by using Log-rank test and one-way ANOVA followed by *post hoc*-Dunnett’s test was used for mean pupation time.

### Statistical analysis

One-way ANOVA or two-way ANOVA followed by *post hoc* Dunnett’s or Tukey’s HSD multiple comparisons were used respectively when data were normally distributed. For data sets that did not have a normal distribution, nonparametric Kruskal–Wallis test followed by Dunn’s *post hoc* multiple comparisons was used. The statistical analyses and sleep data analyses were performed using GraphPad PRISM version 9.2.0. Error bars in all the box-whisker plots represent inter quartile range and the error bars in the waveform of sleep, waveform of activity rest rhythm and percentage pupariation graph represents standard error of the mean (SEM).

## Results

### Short term exposure to caffeine reduces sleep in *Drosophila*

Sleep changes with increasing age in *Drosophila*. As caffeine reduces sleep, we assessed the effect of short exposure to caffeine on sleep during different ages namely 1, 10, 20 and 30 day old flies. Freshly emerged 1 day old flies showed a decrease in sleep only at 1 mg/mL of caffeine concentration when compared to the control (Figure 1A, C). Whereas 10, 20 and 30 day old flies showed decrease in sleep in all the three caffeine concentrations namely 0.5, 0.75 and 1 mg/mL when compared to the control (Figure 1A-C, S1A-B) (Statistical details in Table 1).

**Figure 1:**
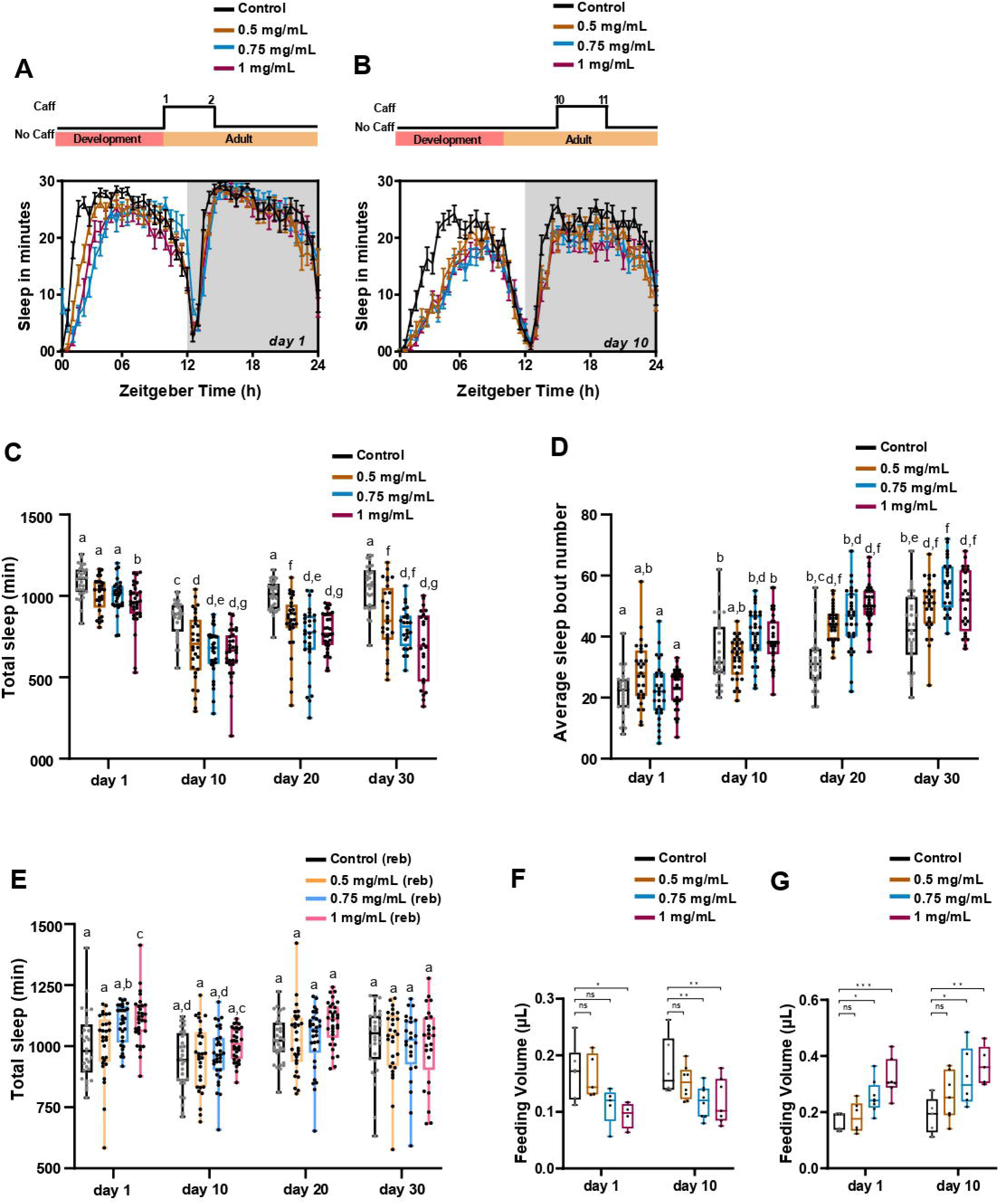
Effect of short exposure to caffeine on sleep and food intake. (A, B) Sleep in min for every 30 min over a period of 24h under LD is shown for both 1 day and 10 day old *w^1118^* flies under short exposure to 0.5 mg/mL, 0.75 mg/mL and 1 mg/mL of caffeine. Schematic on top of the graph depicts short exposure to caffeine. For day 1 old flies the caffeine (Caff) treatment was from day 1 to 2 and for 10 day old flies it was from 10^th^ to 11^th^ day. (C) Quantified total sleep in min for short exposure to caffeine of 1, 10, 20 and 30 day old *w^1118^* flies under 0.5, 0.75 and 1 mg/mL caffeine concentration. Day 1 flies showed decrease in sleep with only 1 mg/mL flies, whereas 10, 20 and 30 day old flies showed decrease in sleep with all caffeine concentrations when compared to the control (Statistical details in Table 1). (D) Average sleep bout number for short exposure to caffeine of 1, 10, 20 and 30 day old *w^1118^* flies. Increased sleep fragmentation was observed in all concentrations of caffeine fed 20 and 30 day old flies when compared to their respective control. (Statistical details in Table 2) (E) Total sleep in min during sleep rebound post short exposure to caffeine of 1, 10, 20 and 30 day old *w^1118^* flies (Statistical details in Table 3). (F) Food intake post 24 hr short exposure to caffeine. Food intake was assessed from ZT 01-04 using CAFE with the presence of caffeine in the food during the assay. Decreased feeding was observed upon presence of caffeine during the CAFE assay (One way ANOVA followed by Dunnett’s multiple comparison Control (day 1) v/s 1 mg/mL (day 1) *p*<0.05) (One way ANOVA followed by Dunnett’s multiple comparison Control (day 10) v/s 0.75 mg/mL (day 10) *p*<0.01; Control (day 10) v/s 1 mg/mL (day 10) *p*<0.001). (G) Food intake post short exposure to caffeine from ZT 01-04 using CAFE without caffeine in the liquid food during the assay. Increased feeding was observed in the absence of caffeine during the CAFE assay (One way ANOVA followed by Dunnett’s multiple comparison Control (day 1) v/s 0.75 mg/mL (day 1) *p*<0.05; Control (day 1) v/s 1 mg/mL (day 1) *p*<0.001) (One way ANOVA followed by Dunnett’s multiple comparison Control (day 10) v/s 0.75 mg/mL (day 10) *p*<0.05; Control (day 10) v/s 1 mg/mL (day 10) *p*<0.01). The same alphabets indicate that they are not statistically different from each other and different alphabets indicate statistically significant differences. The error bars represent the interquartile range.

**Table 1:** Statistical details of effect of short exposure to caffeine on sleep. The table contains statistical details of the 2-way ANOVA followed by Tukey’s post hoc details on short term exposure to caffeine on sleep in day 1, 10, 20 and 30 day old flies. The significance is represented with asterisks (*) and non significant comparisons with ns.

| **Tukey's multiple comparisons test** | **Adjusted *p*-value** |  |
| --- | --- | --- |
| Day 1:Control vs. Day 1:0.5 mg/mL | ns | 0.77 |
| Day 1:Control vs. Day 1:0.75 mg/mL | ns | 0.64 |
| Day 1:Control vs. Day 1:1 mg/mL | * | 0.02 |
| Day 10:Control vs. Day 10:0.5 mg/mL | ** | 0.002 |
| Day 10:Control vs. Day 10:0.75 mg/mL | *** | <0.001 |
| Day 10:Control vs. Day 10:1 mg/mL | *** | <0.001 |
| Day 20:Control vs. Day 20:0.5 mg/mL | ** | 0.001 |
| Day 20:Control vs. Day 20:0.75 mg/mL | *** | <0.001 |
| Day 20:Control vs. Day 20:1 mg/mL | *** | <0.001 |
| Day 30:Control vs. Day 30:0.5 mg/mL | *** | <0.001 |
| Day 30:Control vs. Day 30:0.75 mg/mL | *** | <0.001 |
| Day 30:Control vs. Day 30:1 mg/mL | *** | <0.001 |
| Day 1:Control vs. Day 10:Control | *** | <0.001 |
| Day 1:Control vs. Day 20:Control | ns | 0.66 |
| Day 1:Control vs. Day 30:Control | ns | >0.99 |
| Day 10:Control vs. Day 20:Control | * | 0.02 |
| Day 10:Control vs. Day 30:Control | *** | <0.001 |
| Day 20:Control vs. Day 30:Control | ns | >0.99 |
| Day 1:0.5 mg/mL vs. Day 10:0.5 mg/mL | *** | <0.001 |
| Day 1:0.5 mg/mL vs. Day 20:0.5 mg/mL | ** | 0.001 |
| Day 1:0.5 mg/mL vs. Day 30:0.5 mg/mL | * | 0.04 |
| Day 10:0.5 mg/mL vs. Day 20:0.5 mg/mL | * | 0.03 |
| Day 10:0.5 mg/mL vs. Day 30:0.5 mg/mL | ** | 0.001 |
| Day 20:0.5 mg/mL vs. Day 30:0.5 mg/mL | ns | >0.99 |
| Day 1:0.75 mg/mL vs. Day 10:0.75 mg/mL | *** | <0.001 |
| Day 1:0.75 mg/mL vs. Day 20:0.75 mg/mL | *** | <0.001 |
| Day 1:0.75 mg/mL vs. Day 30:0.75 mg/mL | *** | <0.001 |
| Day 10:0.75 mg/mL vs. Day 20:0.75 mg/mL | ns | 0.79 |
| Day 10:0.75 mg/mL vs. Day 30:0.75 mg/mL | * | 0.04 |
| Day 20:0.75 mg/mL vs. Day 30:0.75 mg/mL | ns | 0.97 |
| Day 1:1 mg/mL vs. Day 10:1 mg/mL | *** | <0.001 |
| Day 1:1 mg/mL vs. Day 20:1 mg/mL | ** | 0.001 |
| Day 1:1 mg/mL vs. Day 30:1 mg/mL | *** | <0.001 |
| Day 10:1 mg/mL vs. Day 20:1 mg/mL | ns | 0.05 |
| Day 10:1 mg/mL vs. Day 30:1 mg/mL | ns | >0.99 |
| Day 20:1 mg/mL vs. Day 30:1 mg/mL | ns | 0.66 |

**Table 2:** Statistical details of effect of short exposure to caffeine on sleep fragmentation. The table contains statistical details of the 2-way ANOVA followed by Tukey’s post hoc details on effect of short term exposure to caffeine on sleep fragmentation in day 1, 10, 20 and 30 day old flies. Rest of the details are the same as Table 1.

| **Tukey's multiple comparisons test** | **Adjusted *p*-value** |  |
| --- | --- | --- |
| Day 1:Control vs. Day 1:0.5 mg/mL | ns | 0.2 |
| Day 1:Control vs. Day 1:0.75 mg/mL | ns | >0.99 |
| Day 1:Control vs. Day 1:1 mg/mL | ns | >0.99 |
| Day 10:Control vs. Day 10:0.5 mg/mL | ns | >0.99 |
| Day 10:Control vs. Day 10:0.75 mg/mL | ns | 0.38 |
| Day 10:Control vs. Day 10:1 mg/mL | ns | 0.82 |
| Day 20:Control vs. Day 20:0.5 mg/mL | *** | <0.001 |
| Day 20:Control vs. Day 20:0.75 mg/mL | *** | <0.001 |
| Day 20:Control vs. Day 20:1 mg/mL | *** | <0.001 |
| Day 30:Control vs. Day 30:0.5 mg/mL | * | 0.04 |
| Day 30:Control vs. Day 30:0.75 mg/mL | *** | <0.001 |
| Day 30:Control vs. Day 30:1 mg/mL | * | 0.01 |
| Day 1:Control vs. Day 10:Control | *** | <0.001 |
| Day 1:Control vs. Day 20:Control | *** | <0.001 |
| Day 1:Control vs. Day 30:Control | *** | <0.001 |
| Day 10:Control vs. Day 20:Control | ns | >0.99 |
| Day 10:Control vs. Day 30:Control | ns | 0.08 |
| Day 20:Control vs. Day 30:Control | *** | <0.001 |
| Day 1:0.5 mg/mL vs. Day 10:0.5 mg/mL | ns | 0.77 |
| Day 1:0.5 mg/mL vs. Day 20:0.5 mg/mL | *** | <0.001 |
| Day 1:0.5 mg/mL vs. Day 30:0.5 mg/mL | *** | <0.001 |
| Day 10:0.5 mg/mL vs. Day 20:0.5 mg/mL | *** | <0.001 |
| Day 10:0.5 mg/mL vs. Day 30:0.5 mg/mL | *** | <0.001 |
| Day 20:0.5 mg/mL vs. Day 30:0.5 mg/mL | ns | 0.48 |
| Day 1:0.75 mg/mL vs. Day 10:0.75 mg/mL | *** | <0.001 |
| Day 1:0.75 mg/mL vs. Day 20:0.75 mg/mL | *** | <0.001 |
| Day 1:0.75 mg/mL vs. Day 30:0.75 mg/mL | *** | <0.001 |
| Day 10:0.75 mg/mL vs. Day 20:0.5 mg/mL | ns | 0.94 |
| Day 10:0.75 mg/mL vs. Day 30:0.75 mg/mL | *** | <0.001 |
| Day 20:0.75 mg/mL vs. Day 30:0.75 mg/mL | *** | <0.001 |
| Day 1:1 mg/mL vs. Day 10:1 mg/mL | *** | <0.001 |
| Day 1:1 mg/mL vs. Day 20:1 mg/mL | *** | <0.001 |
| Day 1:1 mg/mL vs. Day 30:1 mg/mL | *** | <0.001 |
| Day 10:1 mg/mL vs. Day 20:1 mg/mL | *** | <0.001 |
| Day 10:1 mg/mL vs. Day 30:1 mg/mL | *** | <0.001 |
| Day 20:1 mg/mL vs. Day 30:1 mg/mL | ns | >0.99 |

**Table 3:** Statistical details of effect of short exposure to caffeine on rebound sleep. The table contains statistical details of the 2-way ANOVA followed by Tukey’s post hoc details on effect of short term exposure to caffeine on sleep fragmentation in day 1, 10, 20 and 30 day old flies. Rest of the details are the same as Table 1.

| **Tukey's multiple comparisons test** | **Adjusted *p*-value** |  |
| --- | --- | --- |
| Day 1:Control vs. Day 1:0.5 mg/mL | ns | >0.99 |
| Day 1:Control vs. Day 1:0.75 mg/mL | ns | 0.19 |
| Day 1:Control vs. Day 1:1 mg/mL | * | 0.03 |
| Day 10:Control vs. Day 10:0.5 mg/mL | ns | >0.99 |
| Day 10:Control vs. Day 10:0.75 mg/mL | ns | >0.99 |
| Day 10:Control vs. Day 10:1 mg/mL | ns | 0.72 |
| Day 20:Control vs. Day 20:0.5 mg/mL | ns | >0.99 |
| Day 20:Control vs. Day 20:0.75 mg/mL | ns | >0.99 |
| Day 20:Control vs. Day 20:1 mg/mL | ns | 0.69 |
| Day 30:Control vs. Day 30:0.5 mg/mL | ns | >0.99 |
| Day 30:Control vs. Day 30:0.75 mg/mL | ns | >0.99 |
| Day 30:Control vs. Day 30:1 mg/mL | ns | >0.99 |
| Day 1:Control vs. Day 10:Control | ns | 0.88 |
| Day 1:Control vs. Day 20:Control | ns | >0.99 |
| Day 1:Control vs. Day 30:Control | ns | >0.99 |
| Day 10:Control vs. Day 20:Control | ns | 0.24 |
| Day 10:Control vs. Day 30:Control | ns | 0.29 |
| Day 20:Control vs. Day 30:Control | ns | >0.99 |
| Day 1:0.5 mg/mL vs. Day 10:0.5 mg/mL | ns | 0.91 |
| Day 1:0.5 mg/mL vs. Day 20:0.5 mg/mL | ns | >0.99 |
| Day 1:0.5 mg/mL vs. Day 30:0.5 mg/mL | ns | >0.99 |
| Day 10:0.5 mg/mL vs. Day 20:0.5 mg/mL | ns | 0.61 |
| Day 10:0.5 mg/mL vs. Day 30:0.5 mg/mL | ns | 0.82 |
| Day 20:0.5 mg/mL vs. Day 30:0.5 mg/mL | ns | >0.99 |
| Day 1:0.75 mg/mL vs. Day 10:0.75 mg/mL | ** | 0.005 |
| Day 1:0.75 mg/mL vs. Day 20:0.75 mg/mL | ns | 0.88 |
| Day 1:0.75 mg/mL vs. Day 30:0.75 mg/mL | ns | 0.34 |
| Day 10:0.75 mg/mL vs. Day 20:0.75 mg/mL | ns | 0.69 |
| Day 10:0.75 mg/mL vs. Day 30:0.75 mg/mL | ns | >0.99 |
| Day 20:0.5 mg/mL vs. Day 30:0.75 mg/mL | ns | >0.99 |
| Day 1:1 mg/mL vs. Day 10:1 mg/mL | ns | 0.06 |
| Day 1:1 mg/mL vs. Day 20:1 mg/mL | ns | >0.99 |
| Day 1:1 mg/mL vs. Day 30:1 mg/mL | ns | 0.1 |
| Day 10:1 mg/mL vs. Day 20:1 mg/mL | ns | 0.22 |
| Day 10:1 mg/mL vs. Day 30:1 mg/mL | ns | >0.99 |
| Day 20:1 mg/mL vs. Day 30:1 mg/mL | ns | 0.31 |

Further, we observed a significant decrease in sleep from day 1 to day 10 old control flies due to sleep ontogeny, as already reported in previous studies^25,32^ (Figure 1A and C). Although sleep decreases during the first 10 days, sleep increases post ∼15 days of emergence with respect to 10 day old flies^25,32^. Similar changes in sleep were observed in our study as well, where 20 day and 30 day old control flies showed increased sleep when compared to 10 day old control flies (Figure 1C). Furthermore, even with 0.5 mg/mL caffeine a similar increase in sleep from day 10 to 20 was observed, whereas this increase in sleep from 10 to 20 was not observed in higher concentrations namely 0.75 and 1 mg/mL caffeine concentrations. This indicated that higher concentrations of caffeine caused a larger decrease in sleep in older flies (Figure 1C) (Statistical details in Table 1). In addition, there was no significant difference in sleep between 20 and 30 day old caffeine treated flies, when compared to caffeine counterparts, except in 0.75 mg/mL. These results suggest that sleep changes with age and short term exposure to only high caffeine concentration decreases sleep in young flies whereas even lower concentrations of caffeine leads to significant decrease in sleep in older flies (Figure 1C) (Statistical details in Table 1).

### Short term exposure to caffeine increases the sleep fragmentation with age in *Drosophila*

Caffeine consumption increases sleep fragmentation in flies^33^. To assess if fragmentation of sleep due to caffeine is dependent on age, we provided short term exposure of caffeine to different age groups of flies and measured their sleep bout number. More sleep episodes translates to shorter sleep bouts leading to fragmented sleep. Upon assessing sleep bout numbers across the different ages, it was observed that sleep fragmentation is increased in 10, 20 and 30 day old flies when compared to day 1 control flies indicating that sleep fragmentation increases with age (Figure 1D) (Statistical details in Table 2). Further, caffeine consumption did not increase the sleep fragmentation in both 1 and 10 day old flies (Figure 1D). Upon caffeine ingestion 20 and 30 day old flies showed higher sleep fragmentation in 0.5 and 1 mg/mL caffeine concentrations when compared to the control group as well as their counterparts from day 1 and day 10 old flies (Figure 1D) (Table 2). Whereas no significant difference was observed in sleep fragmentation between 10 and 20 day old flies fed with 0.75 mg/mL caffeine. But a significant increase in sleep fragmentation was observed between 20 and 30 day old flies fed with 0.75 mg/mL caffeine. These results indicate that short exposure to caffeine increases sleep fragmentation with increasing age in flies.

### Short term exposure to caffeine does not affect the homeostatic sleep in older flies

To test whether short term exposure to caffeine affects homeostatic sleep, we assessed the sleep rebound of caffeine treated flies under various age groups. An increase in sleep upon removal of caffeine, was observed in freshly emerged flies fed with 1 mg/mL caffeine (Figure 1E, S1C). This indicates that homeostatic sleep pathway could be playing a role in caffeine mediated sleep changes in freshly emerged flies but in older flies as we did not find any sleep rebound after removal of caffeine (Figure 1E, S1C-F). To test this further, we analyzed the transcript level of genes known to be upregulated upon sleep deprivation namely *bruchpilot*, *synaptotagmin*, *syntaxin-18* and *homer*^34,35^ in 10 day old flies treated with 1 mg/mL caffeine for 24h along with the control. Only 1 mg/mL was used for this transcript analysis as there was no significant difference in sleep rebound of 10 day old flies across different caffeine concentrations. Surprisingly, none of these genes were upregulated in caffeine treated day 10 old flies when compared to the control (Figure S1G). Older flies were not tested in this assay as there was no difference in sleep rebound between 10, 20 and 30 day old flies.

### Short term exposure to caffeine affects food intake in flies

The decrease in sleep observed upon caffeine ingestion was not mediated through the homeostatic pathway in older flies. Previously, it has been shown that starvation mediated sleep loss is independent of the homeostatic sleep pathway^36^. As caffeine is bitter to taste we hypothesized that decreased sleep could be observed due to reduced feeding. To analyze this, we assessed the food intake on both day 1 and day 10 old flies after 24h of short term exposure to caffeine. Food intake was recorded in the medium containing the respective caffeine concentrations and we observed that presence of caffeine in the medium caused a reduction in food intake compared to the control flies. In freshly emerged flies only 1 mg/mL flies showed a significant decrease in feeding, coinciding with the observed decrease in sleep (Figure 1F), whereas in mature 10 day old flies a significant decrease in feeding was observed in both 0.75 and 1 mg/mL caffeine concentrations (Figure 1F). Furthermore, feeding increased immediately upon removal of caffeine post 24h of caffeine treatment in both day 1 and day 10 old flies. In both day 1 and day 10 old flies a significant increase in food intake was observed upon removal of 0.75 and 1 mg/mL of caffeine concentration (Figure 1G). Feeding was not measured in 20 and 30 day old flies. These results suggest that short exposure to caffeine reduces food intake in flies.

### Prolonged exposure to caffeine does not affect sleep and food intake in flies

Previously it has been shown that prolonged caffeine ingestion has altered effects when compared to short exposures^37,38^. Furthermore, it has also been previously shown that prolonged exposure causes tolerance to caffeine, leads to drowsiness and also cognitive decline in humans (Reference). As the effect of caffeine is dependent on the duration of caffeine intake, we decided to understand its effects on sleep in *Drosophila*. To assess the effect of prolonged caffeine exposure on sleep, we reared larvae under cornmeal dextrose medium containing different concentrations of caffeine namely 0.5, 0.75 and 1 mg/mL. Caffeine ingestion during the pre-adult development delayed the pupation time (Figure S2A). The developmental delay was quantified and a significant increase in pupation time was observed in 0.75 mg/mL and 1 mg/mL concentrations of caffeine (Figure S2A, B). Although caffeine treatment during pre-adult development increased pupation time, we did not observe any lethality during adult emergence.

Even after the emergence, the freshly emerged flies were reared in their respective caffeine concentrations and the activity-rest rhythm was recorded under caffeine post 10 days of emergence. Surprisingly, upon prolonged exposure to caffeine the flies did not exhibit any significant difference in sleep compared to the control (Figure 2A, B). Although there was no difference in total sleep, there could be an effect on sleep quality. To validate this we estimated the mean sleep bout length in flies with prolonged caffeine treatment. Prolonged caffeine treatment did not affect the sleep depth and quality (Figure S2C). We also assessed the effect of prolonged caffeine treatment on feeding and found that flies under prolonged caffeine treatment did not exhibit any significant difference in food intake compared to the control (Figure 2C). Further, we analyzed the food intake on removal of caffeine post ten days of adult caffeine treatment. No difference in feeding was observed even after the removal of caffeine (Figure S2D). This indicates that flies with prolonged exposure to caffeine probably developed caffeine tolerance and it abolished the effect of caffeine on sleep and food intake.

**Figure 2:**
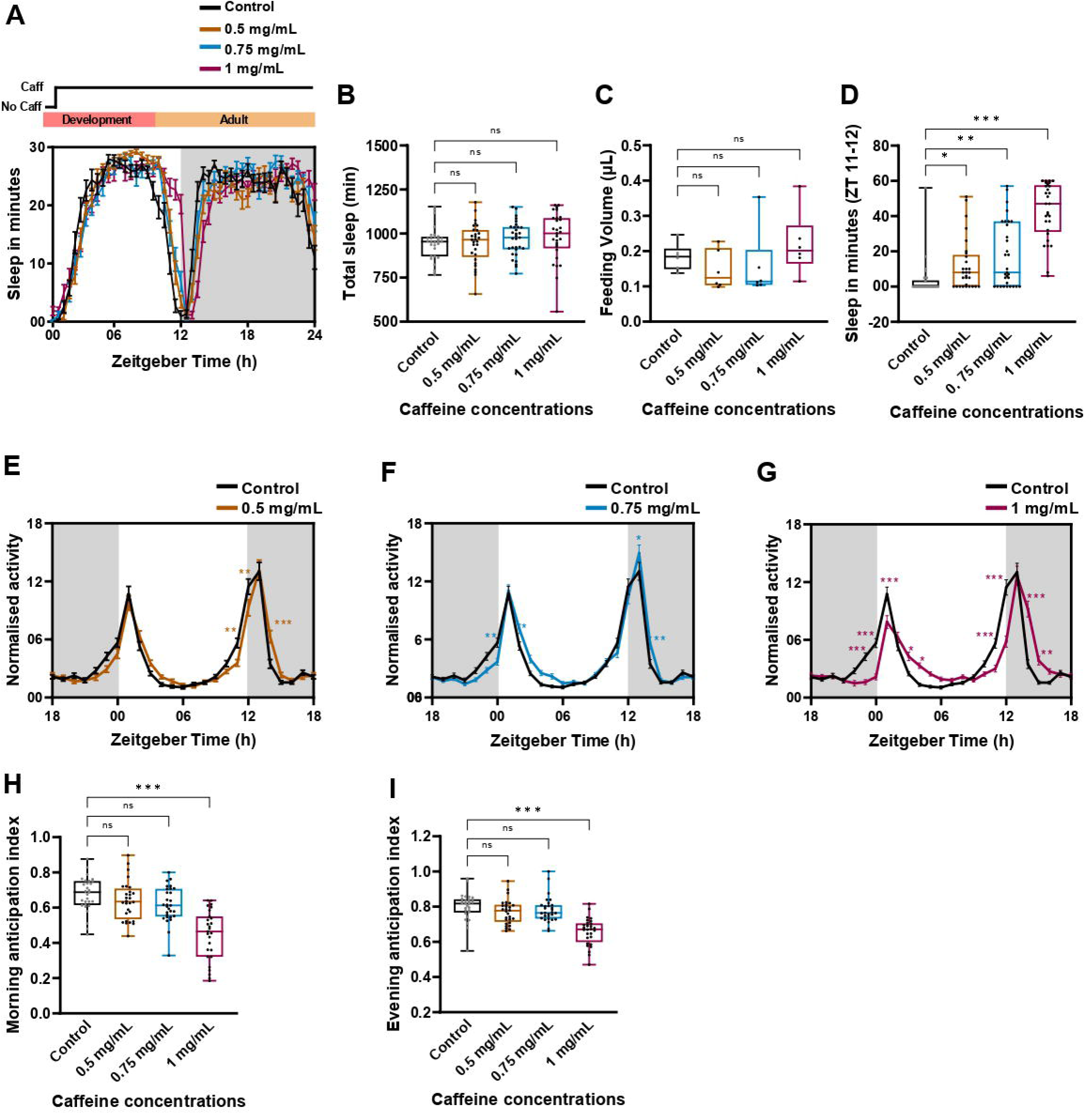
Prolonged caffeine treatment alter morning and evening anticipatory activity. (A) Sleep in min for every 30 min over a period of 24h under LD cycle is shown for 10 day old *w^1118^* flies under 0.5 mg/mL, 0.75 mg/mL and 1 mg/mL of prolonged caffeine treatment. Schematic on top of the graph illustrates the prolonged caffeine treatment protocol. From 1st instar larval stages till the end of the experiment the flies were provided with caffeine containing cornmeal dextrose medium. (B) Total sleep in min for 10 day old *w^1118^* flies under 0.5, 0.75 and 1 mg/mL of prolonged caffeine treatment. (C) Food intake post prolonged caffeine exposure for 10 day old *w^1118^* flies. Food intake was measured at ZT 01-04 using CAFE with the presence of caffeine during the assay. (D) Quantified total sleep in min from ZT 11-12 for prolonged exposure to caffeine for 10 days in *w^1118^* flies under 0.5, 0.75 and 1 mg/mL caffeine concentration. (Kruskal-Wallis test followed by Dunn’s multiple comparisons control v/s 0.5 mg/mL *p*<0.05; control v/s 0.75 mg/mL *p*<0.004; control v/s 1 mg/mL *p*<0.001). (E-G) The percentage of activity after prolonged caffeine treatment for 10 day old flies under 0.5mg/mL, 0.75mg/mL and 1 mg/mL of caffeine concentrations. The percentage of activity with 1h bin-length averaged over 5 consecutive LD is plotted along the *y*-axis and Zeitgeber Time in h along the *x*-axis. The gray shaded area represents the duration of darkness in the LD. (H-I) Morning and evening anticipation index for the flies under 0.5mg/ml, 0.75mg/ml and 1 mg/ml of prolonged caffeine treatment. Flies exhibited a significant reduction in morning and evening anticipation index at 1 mg/mL caffeine concentration when compared to the control. (Kruskal-Wallis test followed by Dunn’s multiple comparisons, morning anticipation index control v/s 1 mg/mL *p*<0.001; evening anticipation index *p*<0.001)

Although flies did not exhibit a significant difference in sleep, a visible delay in siesta offset during the transition from day to night was observed in caffeine-fed flies (Figure 2A). In accordance with that, to understand if caffeine ingestion might cause delay in sleep onset during the transition from day to night, we quantified the sleep latency of caffeine treated flies with respect to the control. We did not find any significant difference in sleep latency (Figure S2E) and we further quantified the sleep during the siesta offset, i.e. ZT 11-12. This indeed showed that flies fed with caffeine had significantly enhanced sleep at ZT 11-12 when compared to the control (Figure 2D). To check if there was a delay in siesta onset we quantified sleep at ZT 03-04. A significant decrease in sleep was observed at 1 mg/mL which indicates that higher caffeine concentration delayed siesta onset as well (Figure S2F). As the timing of siesta onset and offset was altered, this indicated that it could be due to the effect of caffeine on the circadian clock mediated evening anticipatory activity. To investigate this, we recorded the locomotory activity-rest rhythm of flies under prolonged caffeine treatment.

### Prolonged caffeine exposure alters the anticipatory activity and the free running periodicity

As we observed a delay in siesta offset we further examined the effect of prolonged caffeine treatment on activity-rest rhythm of 10 day old flies under LD. Upon 5 days of activity-rest rhythm recording under LD, it was observed that flies fed with 0.5 mg/mL caffeine concentration reduced the activity prior to lights off (Figure 2E) (Statistical details in Table S2). By increasing the caffeine concentration to 0.75 mg/mL, the morning activity prior to lights on was reduced, but no effect on activity prior to lights off was observed (Figure 2F) (Statistical details in Table S2). Whereas by increasing the concentration to 1 mg/mL, activity onset prior to light on and lights off was phase delayed with an increase in the activity after the morning and evening activity peaks (Figure 2G) (Statistical details in Table S2). This change in the activity prior to lights-on and lights-off indicate that there could be a change in the clock mediated morning and evening anticipatory activity. To confirm this we quantified their morning and evening anticipation index. Flies treated with 1 mg/mL of caffeine showed significant decrease in both the morning and evening anticipation index when compared to the control (Figure 2H, 2I).

Further, to test the effect of prolonged caffeine treatment on the endogenous clock, we recorded their activity-rest rhythm under constant darkness (DD). Upon recording the activity-rest rhythm under DD, we observed that prolonged caffeine consumption either lengthened the free running period (Figure 3A-D) or resulted in behavioral arrhythmicity when compared to the control (Figure 3E-G). 93.4% of flies showed rhythmicity in control, whereas percentage rhythmicity decreased with increasing caffeine concentration. 81.8% of flies in 0.5 mg/mL, 81.9% of flies in 0.75 mg/mL and only 37.5% flies in 1 mg/mL of caffeine exhibited rhythmicity (Figure 3H). To further assess the strength of the rhythmicity we analyzed the rhythmicity index. With increasing caffeine concentration more flies showed either weaker rhythmicity or they became arrhythmic (Figure 3I). These results suggest that prolonged caffeine exposure disrupts the free running rhythm in *Drosophila*.

**Figure 3:**
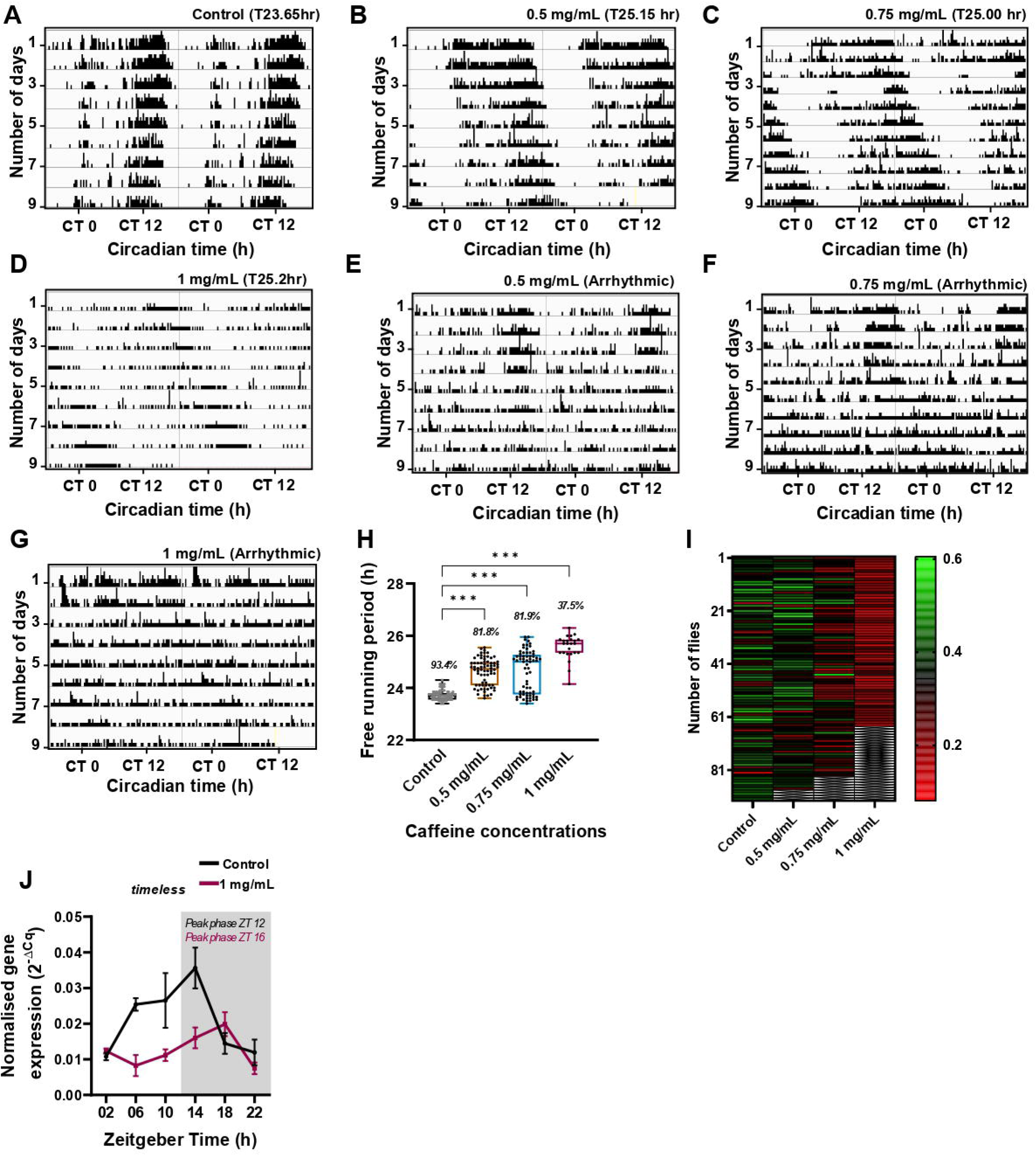
Prolonged caffeine consumption delays free running periodicity and causes arrhythmicity. (A-G) Representative actograms of prolonged caffeine treatment with 0.5, 0.75 and 1 mg/mL caffeine concentration. As the caffeine concentration increases flies exhibit either longer free running periodicity or arrhythmicity under DD. (H) Mean free running periodicity of flies that are rhythmic under different caffeine concentrations. Only flies exhibiting rhythmicity under different caffeine concentrations were selected for this analysis. Numbers given in italics on top of the box-whisker plot represent the percentage of rhythmic flies (One-way ANOVA followed by Dunnett’s multiple comparisons control v/s 0.5 mg/mL *p*<0.001; control v/s 0.75 mg/mL *p*<0.001; control v/s 1mg/mL *p*<0.001). (I) Rhythmicity index of flies under prolonged caffeine treatment with caffeine concentrations ranging from 0.5 mg/mL - 1 mg/mL. Rhythmicity index values >0.3 are rhythmic, and values <0.3 are weakly rhythmic. As the caffeine concentration increases more and more flies show decreased rhythmicity power under DD. The grey criss-cross lines in each vertical bar indicate the dead flies. (J) *timeless* transcript oscillation across 24 hr time-points under LD for control and 1mg/mL of prolonged caffeine treatment. The acrophase (peak phase) of the oscillation was calculated using metacycle analysis. Control showed a peak at ZT 12 and 1 mg/mL of caffeine treated flies showed a peak at ZT 16.

### Prolonged caffeine exposure causes phase delay in the *timeless* transcript oscillation

As we observed a significant difference in behavioral rhythm which is associated with the molecular clock, we assessed the effect of prolonged caffeine treatment on the molecular clock. To this end we measured the mRNA level of *timeless* under LD across a 24h period, with 4h intervals in control and 1 mg/mL caffeine fed flies. Only 1 mg/mL was used for the above assay, because the highest effect was observed with 1 mg/mL caffeine fed flies. The core clock gene *timeless* transcript exhibited a diurnal oscillation with a peak at ZT 12 under LD in control whereas this oscillation was phase delayed by 04h (ZT 16) in 1 mg/mL of prolonged caffeine treatment (Figure 3J) (Table S3). These results indicate that prolonged caffeine treatment affects the molecular clock and causes a phase delay in *timeless* transcript oscillation.

### Effect of prolonged caffeine exposure on circadian clock is largely adult stage specific

To understand whether the effect of prolonged caffeine treatment on circadian clock is development stage mediated or adult stage specific, we provided caffeine treatment only after the flies emerged from the pupae in an adult stage specific manner. After 10 days of adult stage specific caffeine treatment, we recorded their activity rest rhythm with caffeine. In accordance with prolonged caffeine treatment adult stage specific caffeine treatment also delayed the siesta offset when compared to the control. This delayed siesta offset led to increased sleep during ZT 11-12 (Figure 4A, C) and also prior to lights-on (ZT 23-24) (Date not provided). Further, adult stage specific caffeine treatment showed delay in siesta onset which resulted in a decrease in sleep from ZT 03-04 in 0.75 and 1 mg/mL of caffeine concentrations when compared to the control (Figure S3A). Apart from delaying the siesta offset, a significant difference in sleep latency was observed in flies treated with 1 mg/mL caffeine concentration (Figure S3B). We did not observe any significant difference in sleep quality, which was quantified using mean sleep length (Figure S3C). Furthermore, adult stage specific caffeine treatment did not affect the food intake in presence of caffeine (Figure 4D). In addition, adult stage specific caffeine treatment for 10 days post emergence increased total sleep in 0.75 mg/mL and 1 mg/mL caffeine fed flies when compared to the control (Figure 4A-B). This was indeed surprising, as caffeine is known to only decrease sleep or cause tolerance which leads to no effect on sleep.

**Figure 4:**
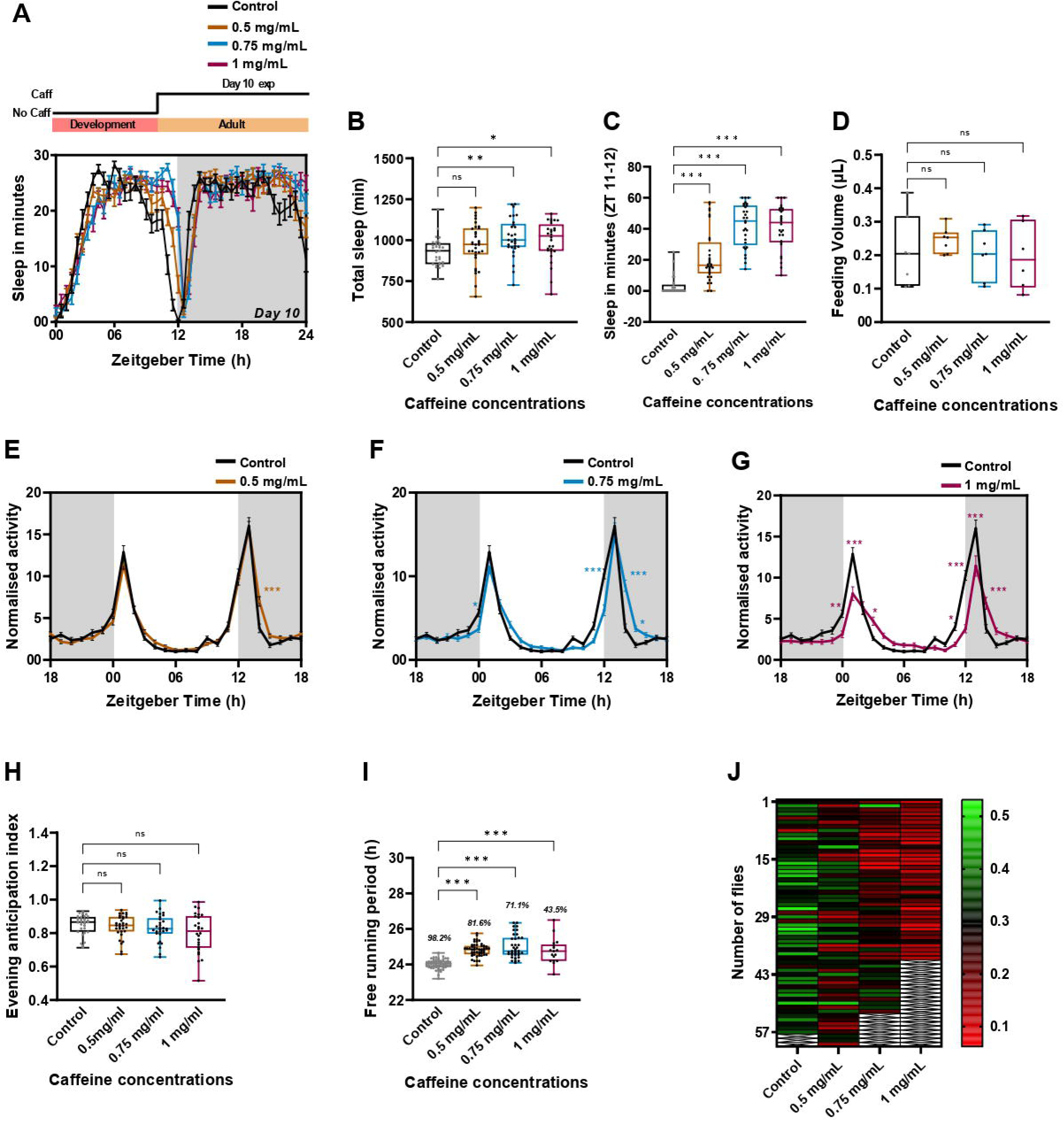
Effect of prolonged caffeine treatment on circadian rhythm is largely adult-stage specific. (A) Sleep in min for every 30 min over a period of 24h under LD for10 day old *w^1118^* flies under adult stage specific caffeine treatment of 0.5, 0.75 and 1 mg/mL of caffeine along with the control. Schematic on top of the graph illustrates adult stage specific caffeine treatment. Flies were fed with caffeine post emergence from day 0 till the end of the experiment. (B) Total sleep in min for *w^1118^* flies under 0.5, 0.75 and 1 mg/mL of adult stage specific caffeine treatment for 10 days. Increased sleep was observed in 0.75 and 1mg/mL caffeine concentration. (Kruskal-Wallis Test followed by Dunn’s multiple comparisons; control v/s 0.75 mg/mL *p*< 0.003; control v/s 1 mg/mL *p*<0.01). (C) Quantified total sleep in min from ZT 11-12 in adult stage caffeine treatment for 10 days in *w^1118^* flies under 0.5, 0.75 and 1 mg/mL caffeine concentration. (Kruskal-Wallis Test followed by Dunn’s multiple comparisons; control v/s 0.5 mg/mL *p*<0.001; control v/s 0.75 mg/mL *p*<0.001; control v/s 1 mg/mL *p*<0.001). (D) Food intake post adult stage caffeine exposure for 10 days from ZT 01-04 using CAFE without caffeine during the assay. (E-G) The percentage activity of 10 days of adult stage caffeine treatment with 0.5, 0.75 and 1 mg/mL caffeine concentration (Statistical details are provided in Table S2). (H) Anticipation to lights-off is plotted for the flies under 0.5 mg/mL, 0.75 mg/mL and 1 mg/mL of prolonged caffeine treatment. No significant difference in anticipation is observed. (I) Mean free running periodicity of flies that are rhythmic under different caffeine concentrations. Only flies exhibiting rhythmicity under different caffeine concentrations were selected for this analysis. Numbers given in italics on top of the box-whisker plot represent the percentage of rhythmic flies. (Kruskal-Wallis Test followed by Dunn’s multiple comparisons; control v/s 0.5 mg/mL *p*<0.001; control v/s 0.75 mg/mL *p*<0.001; control v/s 1 mg/mL *p*<0.001). (J) Rhythmicity index of flies under adult stage specific caffeine treatment with concentrations ranging from 0.5 mg/mL to 1 mg/mL. As the caffeine concentration increases more and more flies show decreased rhythmic power under DD. Rest of the experimental details are the same as in Figure 2, 3.

To further assess the effect of adult-stage specific caffeine treatment on circadian rhythm, we recorded their activity-rest rhythm under LD with caffeine for five consecutive days. In accordance with prolonged caffeine consumption, 0.5 mg/mL of adult stage specific treatment caused an increased activity after the evening activity peak (Figure 4E) (Statistical details in Table S4). Whereas when the concentration was increased to 0.75 mg/mL, apart from observing decreased activity coinciding with lights on and lights off, these flies also exhibited an increased activity after the evening activity peak (Figure 4F) (Statistical details in Table S4). Similar to prolonged caffeine treatment, 1 mg/mL caffeine concentration decreased activity prior to lights on and lights off and increased the activity after the morning and evening activity peaks (Figure 4G) (Statistical details in Table S4). This altered activity during the morning and evening hours was further quantified using the morning and evening anticipation index but no significant difference was observed (Figure 4H).

Further, we also assessed the effect of adult-stage specific caffeine treatment on endogenous clocks by recording their activity under DD for 10 consecutive days and assessed their free running periodicity. Similar to the prolonged caffeine treatment, adult-stage specific caffeine treatment also led to either longer free-running periodicity (Figure 4I) or arrhythmicity (Figure 4J). These results suggest that effects of prolonged caffeine treatment on circadian clock are mostly arising from the adult stage whereas effects on sleep could be influenced from development as well.

### Effect of prolonged caffeine exposure on circadian rhythm is not due to aging

In both the prolonged and adult stage specific caffeine treatment a decrease in lifespan was observed (Figure S3D-E). The average lifespan of a control male fly was close to 75 days with the fifty percent life expectancy (death rate) to be over 60 days. Whereas, in caffeine fed flies this life expectancy decreased drastically with 1 mg/mL caffeine fed flies having the least lifespan (Figure S3D-E). The fifty percent life expectancy of 1 mg/mL fed caffeine flies was close to 25-30 days. The LD and DD activity-rest rhythm recording of flies starting with 10-day old flies took 16 days for completion, by which the age of the fly is close to 25 days i.e. close to the 50% life expectancy of 1 mg/mL caffeine fed flies. Studies in the past have shown that aging dampens the amplitude of circadian rhythm and also leads to longer free running period^39,40^. One of the stark observations we made during prolonged caffeine treatment was longer free running periodicity and arrhythmicity. Therefore, to understand if caffeine mediated premature aging was affecting the circadian rhythm or if caffeine affected the circadian clock, we designed experiments by recording the activity-rest rhythm from a younger age (day 2 old flies, who were fed with different caffeine concentrations post emergence). In this case the age of these flies by the time of completion of activity-rest rhythm recording under LD and DD was only 18 days. By 18 days only less than 10% of fly death was observed in the life span assay of 1 mg/mL caffeine fed flies (Figure S3D-E).

Upon recording activity-rest rhythm with caffeine for 5 consecutive days (day 2-day 7) under LD starting with 2 day old younger flies, it was observed that similar to prolonged caffeine treatment, 0.5 mg/mL showed decreased activity prior to lights off and increased activity post lights off (Figure 5A) (Statistical details in Table S5). By increasing the concentration to 0.75 mg/mL flies still showed a decreased activity prior to lights off and increased activity after lights off without having any effect on the morning peak, as observed in prolonged caffeine treatment (Figure 5B) (Statistical details in Table S5). By increasing the concentration to 1 mg/mL caffeine concentration a decrease in the activity was observed during and before the evening activity peak as well as an increase in the activity after the evening activity peak. But surprisingly under 1 mg/mL caffeine concentration an increase in activity prior to lights on was observed when compared to the control (Figure 5C) (Statistical details in Table S5). Although there was an increase in morning activity prior to lights on, we did not find any significant difference in the morning anticipation index compared to the control (Figure 5D). We further assessed the evening anticipation index and a significant decrease was observed in 1 mg/mL caffeine fed flies when compared to the control (Figure 5E). Following the locomotor activity recording under LD for 5 days, the endogenous free running periodicity was assessed (day 8-day 18). Similar to prolonged caffeine treatment we observed longer free running periodicity in caffeine fed flies when compared to the control (Figure 5F). Further, arrhythmicity increased with increased caffeine concentrations when compared to the control (Figure 5G). These results further indicate that effects of prolonged caffeine treatment on endogenous circadian rhythm is not because of premature aging and is due to the effects of caffeine on circadian clock.

**Figure 5:**
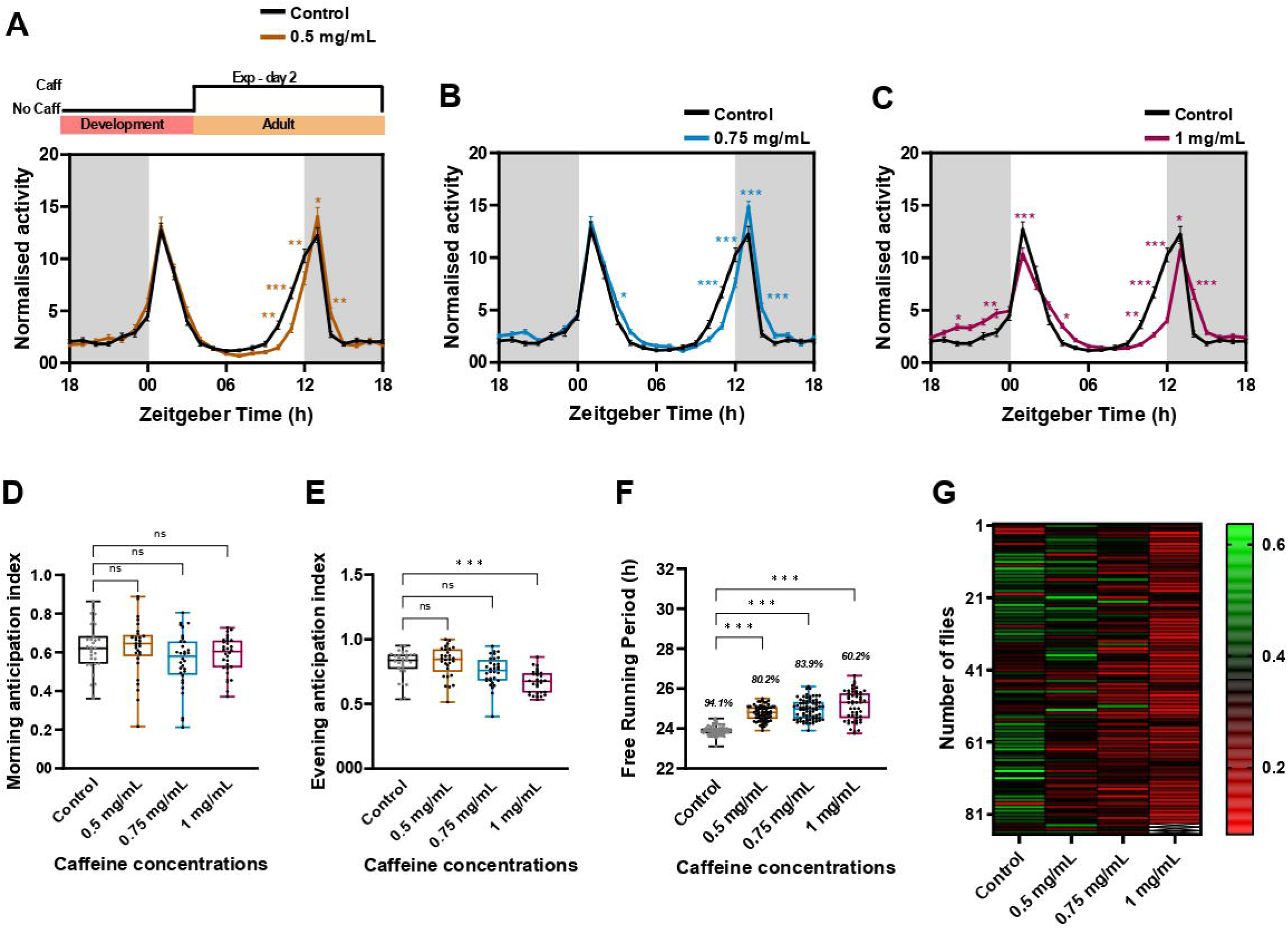
Effect of prolonged caffeine treatment is not because of premature aging. (A-C) The normalized locomotor activity of adult stage specific caffeine treatment under 0.5 mg/mL, 0.75 mg/mL and 1 mg/mL of caffeine concentrations starting with 2 day old flies. Schematic on top of the graph represents the adult stage caffeine treatment. Flies were fed with caffeine post emergence from day 0 till day 2 in vials post which locomotor activity recording under LD with caffeine was initiated. (D, E) Morning and evening anticipation index for the flies under 0.5mg/mL, 0.75 mg/mL and 1 mg/mL of adult specific caffeine treatment. A significant decrease in evening anticipation index was observed in 1 mg/mL caffeine concentration when compared to the control (Kruskal-Wallis Test followed by Dunn’s multiple comparisons; control v/s 1 mg/mL *p*< 0.001). (F) Mean free running periodicity of flies that are rhythmic under different caffeine concentrations. Only flies exhibiting rhythmicity under different caffeine concentrations were selected for this analysis. Numbers given in italics on top of the box-whisker plot represent the percentage of rhythmic flies. (Kruskal-Wallis Test followed by Dunn’s multiple comparisons; control v/s 0.5 mg/mL *p*<0.001; control v/s 0.75 mg/mL *p*<0.001; control v/s 1 mg/mL *p*< 0.001), (G) Rhythmicity index of flies under adult stage specific caffeine treatment in flies in which experiment was started in post 2 days of emergence with concentrations ranging from 0.5 mg/mL-1 mg/mL along with the control. As the caffeine concentration increases more and more flies show decreased rhythmic power. Rest of the experimental details are the same as in Figure 3.

## Discussion

Caffeine, one of the most widely consumed psychostimulants, induces arousal by the activation of dopaminergic 1 receptors (D1R)^24^ and causes sleep fragmentation in flies^24^. Physiological processes are known to deteriorate with age, including the sleep:wake cycle^26,41,42^. With age, sleep consolidation deteriorates by increasing sleep during day-time and also leads to increased wakefulness during night-time, causing shorter sleep episodes^41,42^. Previous studies have already shown that apart from decreasing sleep, caffeine also causes sleep fragmentation^33^. From our study we observe that with increasing age sleep fragmentation increases and this fragmentation is elevated upon caffeine ingestion (Figure 1D). It has been previously shown that sleep loss can lead to accumulation of reactive oxygen species (ROS)^43,44^. With increasing age sleep quality reduces which may also lead to ROS accumalation^42^. Furthermore, treating young flies with paraquet, a ROS inducer increases the sleep fragmentation similar to old flies^42,45^. These results indicate that increase in ROS level leads to sleep fragmentation and vice versa. Furthermore it has been previously shown that cytochrome proteins (CYPs) whose expression levels are known to be repressed by ROS, also had decreased expression upon caffeine ingestion^46^. This indicates that ingestion of caffeine could lead to increased ROS production and that could be implicated in higher sleep fragmentation observed in the present study. Although the molecular basis of sleep fragmentation is yet to be unearthed, our results show that caffeine increases sleep fragmentation with age in flies.

Apart from sleep fragmentation we also observed decreased feeding upon short exposure to caffeine. Furthermore, But, as we see no sleep rebound upon removal of caffeine in older flies, it is evident that decrease in sleep is not mediated by the homeostatic pathway. It has been previously shown that starvation suppresses sleep without a subsequent homeostatic sleep rebound^47^. As we observed a decrease in feeding as a result of short exposure to caffeine, it is likely that reduced feeding may increase the foraging or the starvation mediated hyperactivity^48,49^ and suppress the sleep without inducing a sleep rebound. Reduced feeding indicates that due to its bitter nature, they tend to feed less. Taste sensing neurons or the gustatory receptors Gr93 is known to be involved in sensing caffeine in flies^50^. It has already been shown that absence of these gustatory receptors (Gr93a^3^, _Δ_Gr66a) still shows a decrease in sleep^51,52^. This indicates that the bitter taste to caffeine is not responsible for sleep decrease but the post ingestive effects of caffeine.

From our results and previous experiments it has been shown that effects of caffeine are dependent on the duration of caffeine administration^37,38^. For instance, animals rapidly develop tolerance to caffeine-induced locomotor activity under prolonged caffeine administration^53^. We assessed the effect of prolonged caffeine treatment on flies. 10 days of prolonged caffeine treatment under different concentrations of caffeine did not decrease sleep, but affected the morning, evening anticipatory activity and further led to longer free running period or behavioral arrhythmicity in flies. The effect of prolonged caffeine ingestion on circadian rhythm is not just restricted to flies. It has already been established that caffeine leads to longer free running periodicity in *Neurospora crassa* by increasing the conidiation time. This has been hypothesized to be due to inhibition of cAMP in the cells^54^. Furthermore through in-vivo and in-vitro studies in mice it has been shown that caffeine ingestion causes increased free running period^55,56^. Taken together our results showed short exposure to caffeine reduces sleep whereas prolonged caffeine treatment does not affect sleep and it disrupts the circadian rhythm in flies.

Previously, it has been shown that caffeine induces arousal by acting on the dopaminergic 1 receptors (D1R)^24^, but the effect of caffeine on the molecular circadian clock is not yet well understood. Our study indicates that prolonged caffeine treatment phase delays the *timeless* transcript oscillation. Even though the interaction of caffeine with the circadian clock is yet to be understood, the impact of caffeine on the circadian clock at the molecular level is evident from our studies. Similarly, it has been previously reported that caffeine can phase delay the electrical activity rhythm in the isolated SCN from hamster and rat brain^57,58^. Further studies are required to unravel the underlying mechanisms through which caffeine phase delays the *timeless* transcript oscillation. Nevertheless we cannot exclude the possibility that caffeine acts on additional relevant targets to affect the circadian timing in *Drosophila*. Previous reports showed that caffeine ingestion causes inhibition of cyclic Adenosine Monophosphate Phosphodiesterase (cAMP PDE) activity, through secondary messenger molecule cAMP and activation of ryanodine receptors^33,59^. Caffeine causes widespread elevation of cAMP throughout the brain^33^ Furthermore, it is also shown that cAMP downstream signaling activation by calcium can directly modulate the speed of the circadian clock independent of the TTFL’s^60^. Whether prolonged caffeine treatment affects the phase of the circadian clock through the changes in cAMP level requires further empirical attention. Moreover, the effect of caffeine on behavioral output is dependent not only on the duration of caffeine intake but also on the concentration of caffeine. Higher concentrations of caffeine have shown to cause longer free running periodicity when provided for a small duration^38,56^.

Apart from this our study also showed that prolonged caffeine treatment during larval stages, delays the pre-adult development in *Drosophila*. From previous reports it is known that this caffeine mediated developmental delay is mostly associated with the changes in the expression of Adenosine receptor (dAdoR) that orchestrates development in *Drosophila*^61^. Subsequently, it has been shown that the effect of short exposure to caffeine on sleep is not mediated through adenosine receptor signaling in *Drosophila*^33,62^ unlike mammals^63^. But the impact of caffeine treatment during larval stages on adenosine receptors could have an effect on the adult stage sleep needs to be further investigated. Along with decreased locomotor activity and delayed pre-adult development, prolonged caffeine treatment also reduced life span in flies. Reduction in life span is majorly associated with increased ROS production^64^. As it has been previously shown that caffeine reduces the expression of genes involved in clearing ROS^46^, prolonged caffeine treatment may enhance ROS level leading to reduced life span.

The prolonged caffeine treatment disrupted the circadian clock and a similar phase delay in activity onset under LD, longer free running period and behavioral arrhythmicity was observed when caffeine was provided in an adult stage specific manner. Although, adult stage specific caffeine treatment phase delay the circadian clock, an increase in sleep duration was also observed. The increase in sleep observed is quite surprising as caffeine has never been shown to cause an increase in sleep (Figure B). In a previous study, adult stage caffeine treatment (defined as chronic caffeine treatment in Wu et al., 2009) for 7 days still decreased sleep in flies. But, in our study we did not observe this. One of the major reasons associated with this is the use of a different genotype *RC1* in the previously reported study and *w^1118^* flies were used in our study^33^. RC1 is found to be caffeine sensitive and a decrease in sleep has been observed whereas 10 days of adult stage specific caffeine treatment in *w^1118^* flies increased the sleep. More experimental evidence is required to decode the differential effect of caffeine treatment duration on sleep. Another interesting observation from our study is that the concentration and duration of caffeine treatment plays a critical role in affecting the circadian clock.

In summary, the results of our studies showed that short exposure to caffeine reduces sleep in flies and increases sleep fragmentation with age. While prolonged caffeine treatment does not have an effect on sleep, it affects the morning and evening anticipatory activity in flies. In addition, prolonged caffeine treatment phase delays the transcript oscillation of the clock gene *timeless* and disrupts the circadian rhythm in *Drosophila*.

## Acknowledgements

We thank Dr. Jishy Varghese for the valuable suggestions. This work was supported by the DBT/Wellcome Trust India Alliance Fellowship [IA/E/15/1/502329] awarded to NNK and intramural fund from Indian Institute of Science Education and Research, Thiruvananthapuram.

---

**Figure S1: Effect of short exposure to caffeine on homeostatic sleep.**

(A, B) Sleep in min for every 30 min over a period of 24h under LD is shown for 20 and 30 day old *w^1118^* flies of control, 0.5 mg/mL, 0.75 mg/mL and 1 mg/mL under short exposure to caffeine. Schematic on top of the graph depicts short exposure to caffeine. For day 20 old flies the caffeine treatment was from day 20^th^ to 21^th^ and for 30 day old flies it was from 30^th^ to 31^sh^ day. (C-F) Sleep rebound in min for every 30 min over a period of 24h under LD is shown for 1, 10, 20 and 30 day old *w^1118^* flies immediately after for 24h of short exposure to 0.5 mg/mL, 0.75 mg/mL and 1 mg/mL of caffeine. (G) Quantified mRNA level for *bruchpilot*, *synaptotagmin*, *syntaxin-18* and *homer* of 10 day old flies under short term caffeine exposure to 1 mg/mL concentration.

**Figure S2: Effect of prolonged exposure to caffeine on sleep and development.**

(A-B) Percentage of pupation and mean pupation time in hr under different concentrations of prolonged caffeine treatment. Schematic given above the graph illustrates prolonged caffeine treatment during the larval stages. (C) Mean sleep bout length of 10 day old flies under 0.5, 0.75 and 1 mg/mL caffeine concentrations of prolonged caffeine treatment. No significant difference was observed in sleep bout length when compared to the control. Schematic on top of the graph illustrates the prolonged caffeine treatment protocol. From 1st instar larval stages till the end of the experiment the flies were provided with caffeine containing cornmeal dextrose medium. (D) Food intake post prolonged caffeine exposure for 10 day old flies. Food intake was assessed from ZT 01-04 using CAFE in the absence of caffeine during the assay. No significant difference was observed in food intake when compared to the control. (E) Sleep latency of 10 day old flies with 0.5mg/mL, 0.75 mg/mL and 1 mg/mL of prolonged caffeine treatment. No significant difference was observed in sleep latency when compared to the control. (F) Quantified total sleep in min from ZT 03-04 for 10 day old *w^1118^* flies under 0.5, 0.75 and 1 mg/mL of prolonged caffeine exposure (Kruskal-Wallis Test followed by Dunn’s multiple comparisons control v/s 1 mg/mL *p*< 0.05).

**Figure S3: Effect of adult stage specific caffeine treatment on sleep.**

(A) Quantified total sleep in min from ZT 03-04 with adult stage specific caffeine treatment for 10 days in *w^1118^* flies under 0.5, 0.75 and 1 mg/mL caffeine concentration (Kruskal-Wallis Test followed by Dunn’s multiple comparisons; control v/s 0.75 mg/mL *p*<0.001; control v/s 1 mg/mL *p*< 0.001). (B) Sleep latency of day 10 old flies with 0.5mg/mL, 0.75 mg/mL and 1 mg/mL caffeine concentration under adult stage specific caffeine treatment. 1 mg/mL caffeine concentration showed a decrease in sleep latency when compared to the control (Kruskal-Wallis Test followed by Dunn’s multiple comparisons control v/s 1 mg/mL *p*< 0.05). (C) Mean sleep bout length of 10 day old flies under 0.5, 0.75 and 1 mg/mL adult-stage specific caffeine treatment. No significant difference was observed in sleep bout length when compared to the control. (D) Lifespan of *w^1118^* flies under different concentrations of prolonged caffeine treatment. Percentage of flies alive is plotted on the *y-axis* and time in days is plotted on the *x-axis*. It is represented with the schematics for prolonged caffeine treatment. (E) Lifespan of *w^1118^* flies under different concentrations of adult stage specific caffeine treatment. Percentage of flies alive is plotted on the *y-axis* and time in days is plotted on the *x-axis*. It is represented with the schematics for adult stage caffeine treatment.

**Figure S4: Detailed description of feeding assay.**

(A): Schematic representation of the modified feeding assay using microtips. The 50 ml centrifuge tubes were used for the assay, where a 200 µL tip was inserted into the lid through a hole. A 10 - 20 µL range micro tip was inserted into the 200 µL tip in which 10 µL sucrose solution was provided (with or without caffeine). After the assay, the volume of food intake was calculated by measuring the height of the remaining food volume using a vernier caliper. (B): Food intake of *w^1118^* was measured from ZT 01-04 simultaneously using microcapillary tubes (C) (1 - 5µL range) and microtips (M). No significant difference in food intake volume was observed between the two methods thus verifying our modified feeding assay.

**Table S1: List of primer used for quantitative RT PCR.**

**Table S2: Statistical details of effect of prolonged caffeine treatment on activity rest rhythm.**

The table provides details of Zeitgeber Time (h) where a significant difference in activity rest rhythm was observed in each of the caffeine concentrations(0.5, 0.75, 1 mg/mL) when compared to the control.

**Table S3: Metacycle outcome for *timeless* transcript oscillation.**

The table contains the outcome of JTK analysis performed using metacycle meta2d algorithm followed by combined metacyle analysis. It also provides the *p*-value for the *timeless* transcript in control and 1 mg/mL which defines the oscillation of the transcript. Further, it also contains the phase and period of the transcript under each condition.

**Table S4: Statistical details of effect of adult stage specific caffeine treatment on activity rest rhythm.**

The table provides details of Zeitgeber Time (h) where a significant difference in activity rest rhythm was observed in each of the caffeine concentrations(0.5, 0.75, 1 mg/mL) when compared to the control.

**Table S5: Statistical details of effect of adult stage caffeine treatment on young flies on activity rest rhythm.**

The table provides details of Zeitgeber Time (h) where a significant difference in activity rest rhythm was observed in each of the caffeine concentrations(0.5, 0.75, 1 mg/mL) when compared to the control.

