## Supplementary figures and images for "The duration of caffeine treatment plays an essential role in its effect on sleep and circadian rhythm"

### Figure S1

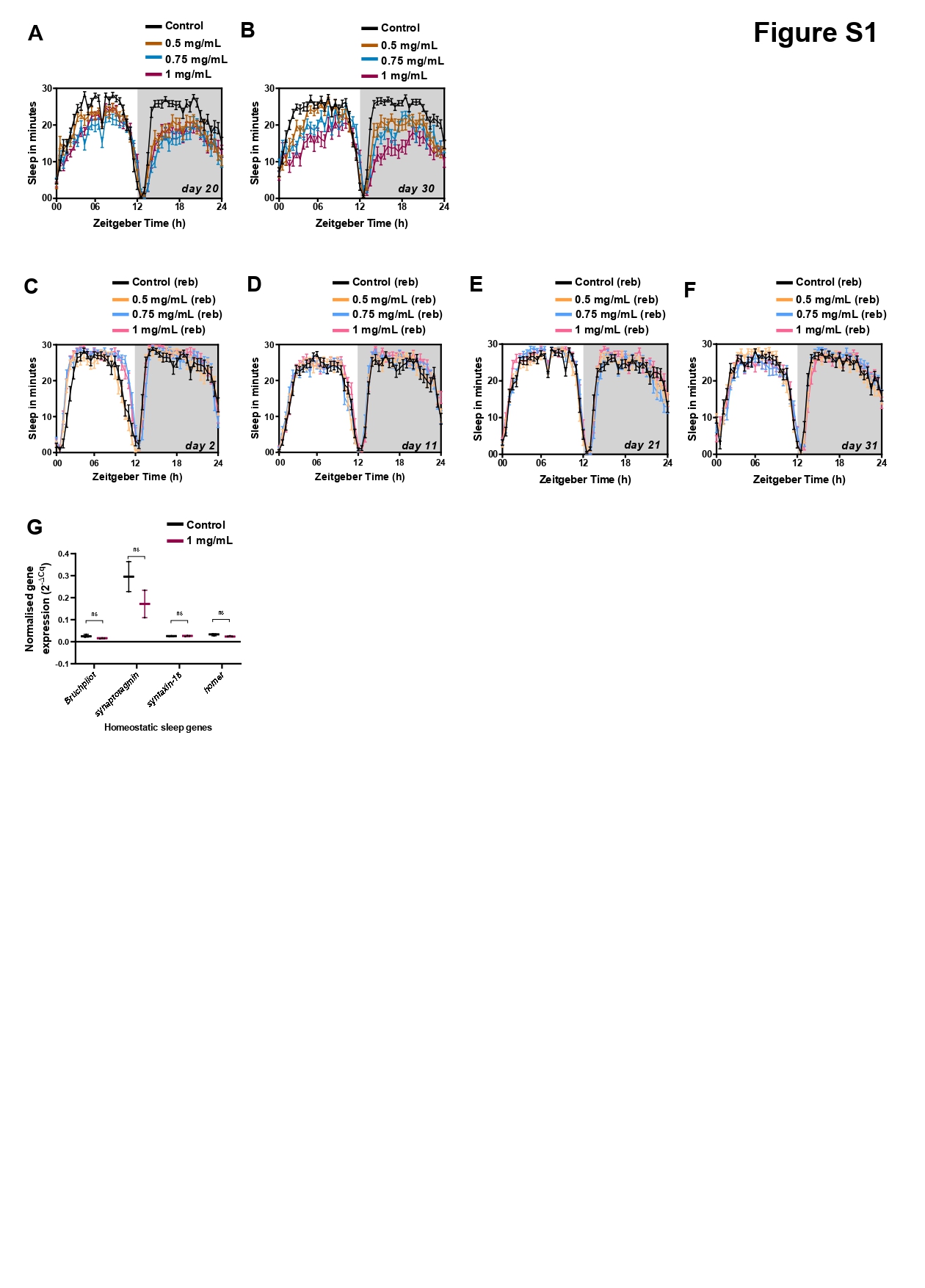

### Figure S2

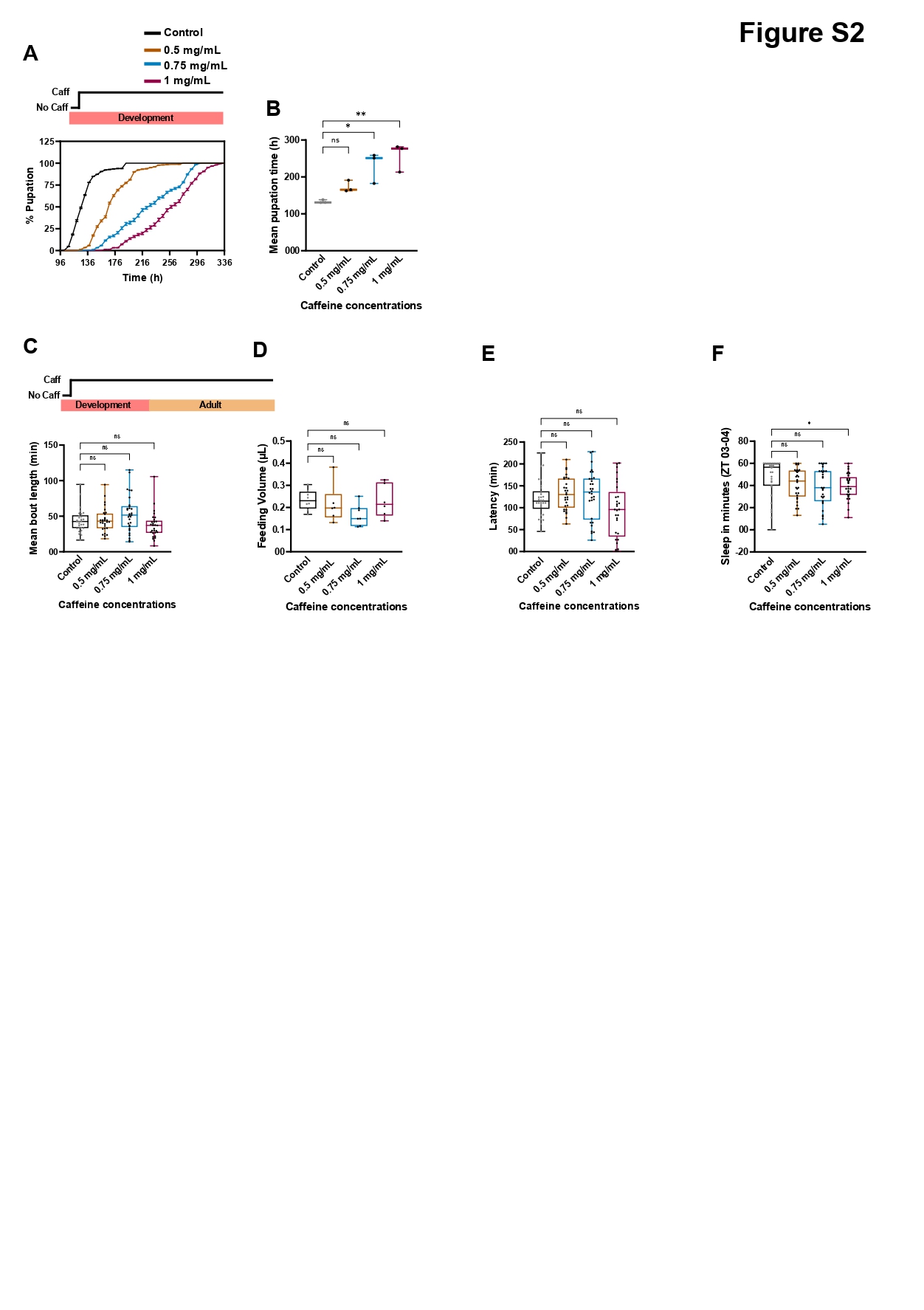

### Figures S3

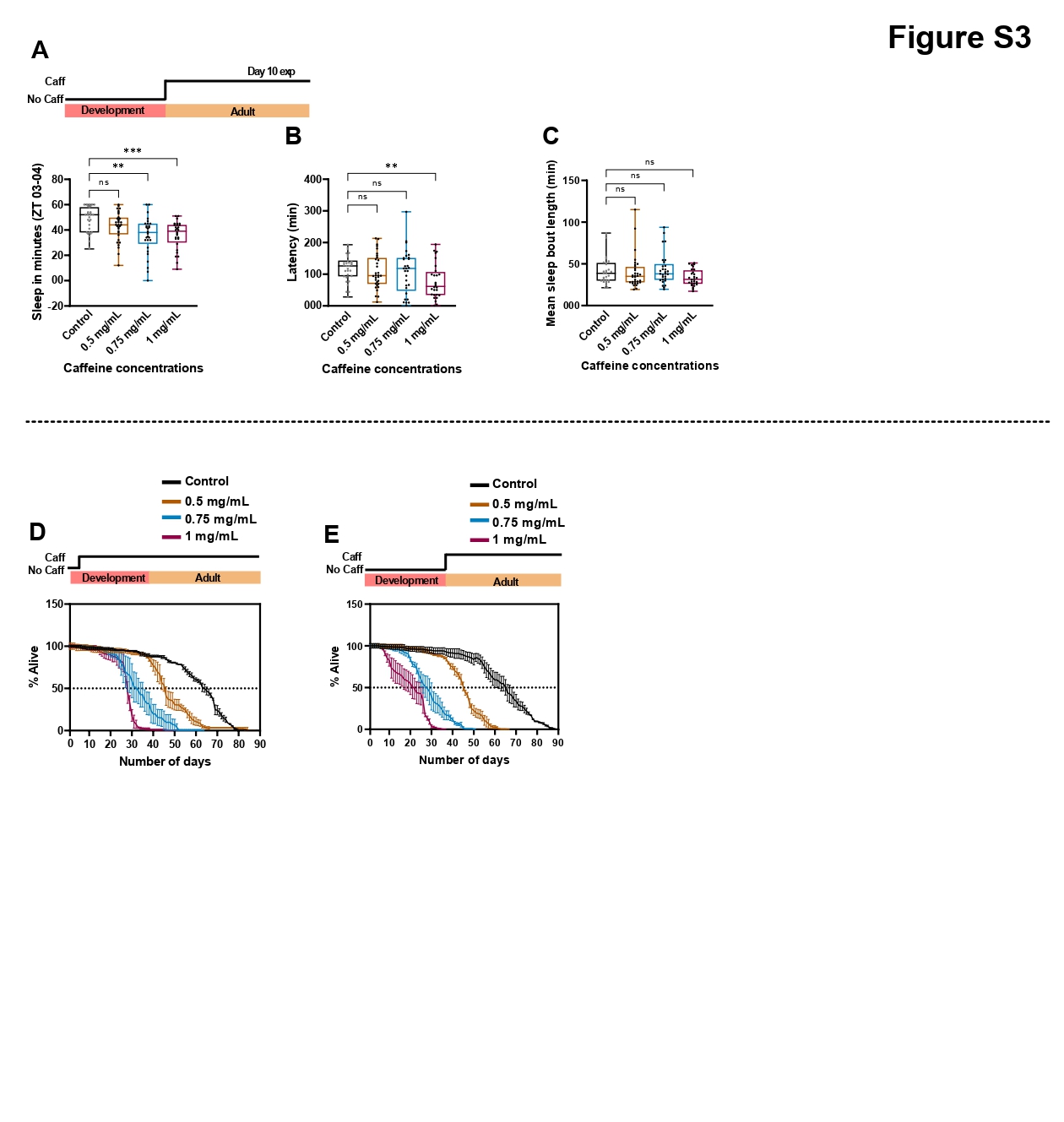
