## Supplementary material for "The duration of caffeine treatment plays an essential role in its effect on sleep and circadian rhythm": Table S1

**Table S1: List of primers used**

| Gene | Forward Primer | Reverse Primer |
|---|---|---|
| *rp49* | GCTAAGCTGTCGCACAAA | TCCGGTGGGCAGCATGTG |
| *timeless* | CCGTGGACGTGATGTACCGCAC | CGCAATGGGCATGCGTCTCTG |
| *homer* | GAACAACCGATTTTCACTGC | GAGCTGTCGTAGAAGGAAGCTAAC |
| *synaptotagmin* | CTGAGTCCGGTCTTCAACGAG | ACACGAGCGTCTTGTTCATGG |
| *bruchpilot* | GCAGTCCATACTACCGCGAC | TTGGATAGTCCATGGCATGGG |
| *Syntaxin 18* | GTGTGCCAACAAGATCACTCGC | GCGTGTAAAAGGTGACGAACTTC |
