## Supplementary material for "The duration of caffeine treatment plays an essential role in its effect on sleep and circadian rhythm": Table S3

### Table S3: Meta2D output results

| | JTK_pvalue | JTK_BH.Q | JTK_period | JTK_adjphase | Meta2d_Base | Meta2d_AMP |
|---|---|---|---|---|---|---|
| control | 0.000115 | 0.000230 | 20 | 12 | 0.0098117 | 0.00932 |
| 1 mg/mL | 0.00754 | 0.00454 | 24 | 16 | 0.00327 | 0.00699 |

The table contains the meta2D output results of JTK algorithm.
