## Supplementary material for "The duration of caffeine treatment plays an essential role in its effect on sleep and circadian rhythm": Table S4

**Table S4:** Effect of adult-stage specific caffeine treatment on activity- rest rhythm of Day 10 old flies.

| **Caffeine concentration: 0.5 mg/mL (comparison with control)** | | |
|---|---|---|
| **Zeitgeber Time (h)** | **Summary** | **Adjusted *p* Value** |
| ZT 14 | *** | <0.001 |
| **Caffeine concentration: 0.75 mg/mL (comparison with control)** | | |
| **Zeitgeber Time (h)** | **Summary** | **Adjusted *p* Value** |
| ZT 00 | * | <0.02 |
| ZT 12 | *** | <0.001 |
| ZT 14 | *** | <0.001 |
| ZT 15 | * | <0.02 |
| **Caffeine concentration: 1 mg/mL (comparison with control)** | | |
| **Zeitgeber Time (h)** | **Summary** | **Adjusted *p* Value** |
| ZT 24 | ** | <0.001 |
| ZT 01 | *** | <0.001 |
| ZT 03 | * | <0.01 |
| ZT 11 | * | <0.9 |
| ZT 12 | *** | <0.001 |
| ZT 13 | *** | <0.001 |
| ZT 14 | *** | <0.001 |

Two-way ANOVA followed by Tukey's post hoc HSD showing Zeitgeber time points (h) at which 0.5, 0.75 and 1 mg/mL caffeine treated flies showed significant difference in activity rest rhythm when compared to the control.
