## Supplementary material for "The duration of caffeine treatment plays an essential role in its effect on sleep and circadian rhythm": Table S5

**Table S5:** Effect of adult- stage specific treatment on activity-rest rhythm of Day 2 old flies.

| **Caffeine concentration: 0.5 mg/mL (comparison with control)** | | |
|---|---|---|
| **Zeitgeber Time (h)** | **Summary** | **Adjusted *p* Value** |
| ZT 10 | ** | <0.002 |
| ZT 11 | *** | <0.001 |
| ZT 12 | ** | <0.002 |
| ZT 13 | * | <0.03 |
| ZT 14 | ** | <0.006 |
| **Caffeine concentration: 0.75 mg/mL (comparison with control)** | | |
| **Zeitgeber Time (h)** | **Summary** | **Adjusted *p* Value** |
| ZT 03 | * | <0.03 |
| ZT 11 | *** | <0.001 |
| ZT 12 | *** | <0.001 |
| ZT 13 | *** | <0.001 |
| ZT 14 | *** | <0.001 |
| **Caffeine concentration: 1 mg/mL (comparison with control)** | | |
| **Zeitgeber Time (h)** | **Summary** | **Adjusted *p* Value** |
| ZT 20 | * | <0.03 |
| ZT 23 | ** | <0.003 |
| ZT 01 | *** | <0.001 |
| ZT 04 | * | <0.01 |
| ZT 10 | ** | <0.002 |
| ZT 11 | *** | <0.001 |
| ZT 12 | *** | <0.001 |
| ZT 13 | * | <0.03 |
| ZT 14 | *** | <0.001 |

Two-way ANOVA followed by Tukey's post hoc HSD showing Zeitgeber time points (h) at which 0.5, 0.75 and 1 mg/mL caffeine treated flies showed significant difference in activity rest rhythm when compared to the control.
